# Differential Effects of Afferent and Efferent Vagus Nerve Stimulation on Gastric Motility Assessed with Magnetic Resonance Imaging

**DOI:** 10.1101/650234

**Authors:** Kun-Han Lu, Jiayue Cao, Robert Phillips, Terry L Powley, Zhongming Liu

## Abstract

**Background:** Vagus nerve stimulation (VNS) is an emerging bioelectronic therapy for regulating food intake and controlling gastric motility. However, the functional impact of different VNS parameters on postprandial gastric motility remains incompletely characterized. Moreover, while most studies focused on stimulating the motor limb (i.e. efferent VNS) of the vagovagal circuitry, the contribution of electrically activating the reflex arc of the circuitry (i.e. afferent VNS) to downstream control of gastric function has seldom been investigated.

**Methods:** Here, we used dynamic gastric magnetic resonance imaging (MRI) to assess antropyloric motility in anesthetized rats during which VNS was applied to the left cervical vagus. The configuration of VNS was varied in terms of directionality (i.e. afferent, efferent or combined afferent and efferent VNS) as well as parameter settings (i.e. pulse amplitude, pulse width, and frequency). The motility measurements were computed using a previously developed computer-assisted image processing pipeline.

**Key Results:** We found that electrical activation that favored the afferent pathway could promote gastric motility and coordination more effectively than direct activation of the efferent pathway. A reduction in antral contraction amplitude and pyloric tone under high-dose efferent VNS highlighted the inhibitory role of the efferent vagovagal circuitry.

**Conclusions & Inferences:** This study demonstrated the direct and reflex gastric responses to cervical VNS measured with MRI. Our findings suggest that selective stimulation of vagal afferents is potentially more favorable than stimulation of vagal efferents in facilitating coordinated antropyloric motility.

**Key Points:**

1. Vagus nerve stimulation is emerging as a new bioelectronic therapy for remedying gastric symptoms. However, the effects of graded VNS settings and directionality preferences of VNS on gastric functions remain incompletely characterized.
2. Dynamic gastric MRI revealed that electrical activation of the afferent pathway could promote antropyloric motility more effectively than direct activation of the efferent pathway.
3. MRI can noninvasively characterize post-prandial gastric motility with high spatial and temporal resolution that could be used to guide the selection of VNS settings.

## Introduction

The parasympathetic innervation to the gastrointestinal (GI) tract plays a key role in regulating, modulating, and controlling GI motility and maintaining energy homeostasis^1,2^. As such innervation is predominantly supplied by the vagus nerve^3,4^, it has been of great interest to modulate motor neural signaling to the gut via electrical stimulation of the vagus nerve^5–7^. However, several morphological and functional studies have suggested that the vagus is a heterogeneous nerve consisting of distinct fiber calibers that carries both efferent and afferent traffic^8,9^. The stimulus-response relations often vary according to the site of stimulation and also the choice of stimulus parameters. As a result, the complete extent of vagal innervation to the enteric neural plexuses remains incompletely characterized.

The vagus is comprised of about 75% sensory afferent and 25% motor efferent fibers^8^. Previous studies have uncovered that, depending on the choice of stimulus parameters, electrical stimulation of efferent fibers could impose both excitatory and relaxatory effect on gastric tone and motility^10–13^. Because of this mixed, opposite effect, it remains unclear whether stimulating the efferent vagal fibers could result in choreographic gastric motility that can be used to remedy gastric symptoms. While a majority of VNS-gastric studies attempted to program the motor limb (i.e. efferent) of the vagovagal circuitry, the effect of stimulating the reflex arc (i.e. afferent) of the circuitry on gut motility is less understood. The significance of afferent stimulation could be evidenced by the fact that reflex vagal excitation by intra-gastric distension has been shown to promote antral motility^14^. Moreover, patients with epilepsy or depression treated with chronic VNS experienced substantial weight loss^15,16^. The influences of VNS on the central nervous system (CNS) may therefore affect appetite control^17,18^, food intake^19^, or potentially lead to top-down modulation on gastric tone and motility. As such, elucidating the potentially differential impact of efferent versus afferent VNS on gastric physiology is necessary for better calibration of neuromodulation efficacy.

To evaluate the efficacy and stability of stimulus settings, most studies employed invasive methods in fasted animals. However, it is not unlikely that the stimulus-response relations might be confounded by such unphysiological assessments. Moreover, as electroceutical therapeutic outcomes in humans are largely derived during meal digestion, the importance of characterizing the stimulation effect under postprandial states should be underscored. In this regard, we have recently demonstrated the use of contrast-enhanced magnetic resonance imaging (MRI) to noninvasively monitor gastric emptying and motility in dietgel-fed rats^20^. The imaging capability has also enabled us to validate the efficacy of left cervical VNS for promoting gastric emptying^21^.

Here, we sought to disentangle the impact of left cervical VNS with different directionality preferences and parameters (i.e. pulse amplitude, pulse width and pulse frequency) on antropyloric motility in anesthetized rats. Dynamic gastric MRI was performed continuously before and during VNS. The occlusive amplitude of antral contraction waves and the luminal size of pyloric opening were measured as a function of stimulus settings. The findings from this study could shed light on the selection of VNS settings for better modulating gastric functions.

## Materials and Methods

### Subjects

Thirteen Sprague-Dawley Rats (Male; Envigo RMS, Indianapolis, IN), ranging from 266 to 338g body weight were used in this study. All experimental procedures were approved by the Purdue Animal Care and Use Committee (PACUC) and the Laboratory Animal Program (LAP). The animals were housed individually in ventilated cages under a strictly controlled environment (temperature:70±2°F, and 12 h light-dark cycle: lights on at 6:00 AM, lights off at 6:00 PM). The floor was elevated by a stainless-steel wire frame during all time to prevent the animals from accessing their feces. All experiments were performed acutely, after which the animals were euthanized according to a standard approved protocol.

### Test Meal

All animals were trained to voluntarily consume a fixed quantity (about 10g) of nutrient-fortified water gel (DietGel Recovery, ClearH2O, ME, USA) under a re-feeding condition following an overnight (18 hours, 5PM to 11AM) fast. The training protocol consisted of 2 stages. During the first stage, an aliquot (~10g) of the Dietgel was placed in a cup in animal’s home cage overnight for 2 times, with the regular rat chow being supplied ad libitum. Once the animal got used to the test meal, their food was deprived for 18 hours followed by re-feeding of the Dietgel. At the end of the 7-day training, all animals were compliant to the naturalistic feeding paradigm in order for us to investigate gastric functions in a physiologically fed condition.

### Animal Preparation and Surgical Implantation of Stimulating Electrode

On the day of imaging, the animal was fed with 10g of Gadolinium (Gd)-labeled test meal after the 18-hour overnight fast. The Gd-labeled test meal was made of 10g Dietgel mixed with 22.4mg Gd-DTPA powder (#381667, Sigma Aldrich, St. Louis, USA) using a double-boiled liquefying approach. Soon after ingestion, the animal was anesthetized with Isoflurane followed by implantation of a bipolar cuff electrode (Pt/Ir electrode; MicroProbes, Gaithersburg, MD, USA) onto the left cervical vagus nerve with a surgical procedure identical to that described in our previous work^21^. The specification of the bipolar cuff electrode is illustrated in Fig. 1A. The animal was then setup in prone position in the scanner. Respiration and body temperature were monitored with an MR-compatible system (SA Instruments Inc., Stony Brook, NY, USA) to ensure stable physiology. The leads of the electrode were connected to a pair of twisted wires that were connected to a current stimulator (A-M Systems Model 2200; A-M Systems, Sequim, WA, USA) placed outside of the MRI room. A bolus injection of 0.01 mg kg^−1^ dexmedetomidine solution (0.05 mg mL^−1^, Zoetis, NJ, USA) was administered subcutaneously (SC). Fifteen minutes after the bolus, dexmedetomidine was infused (SC) continuously throughout the experiment (0.03 mg kg^−1^ h^−1^). The dose of Isoflurane was lowered to 0.3-0.5% as soon as the animal’s respiratory rate began to decrease.

**Figure 1:**
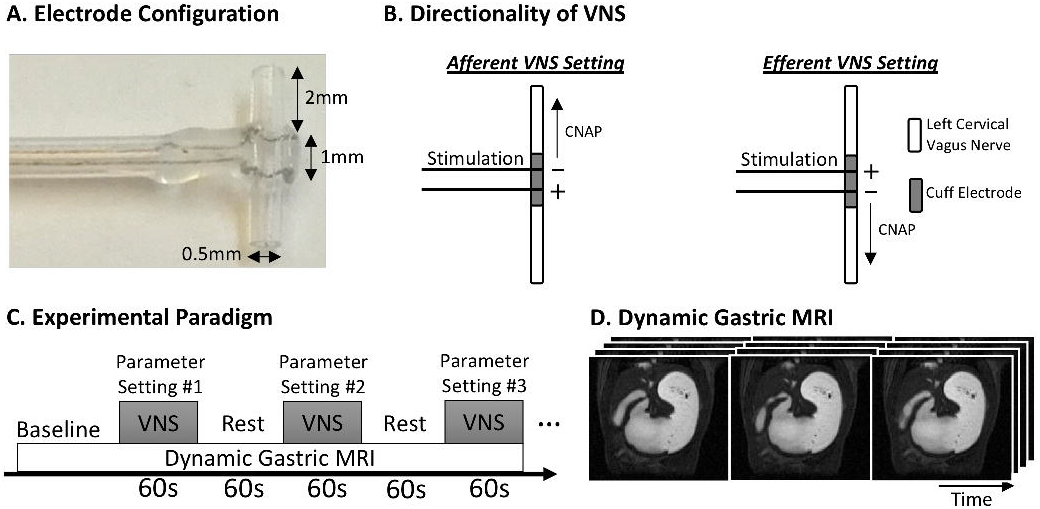
Illustration of instrumentation and experimental design. **(A)** The specification of the bipolar cuff electrode. **(B)** The propagating direction of stimulation-evoked compound nerve action potential was controlled by the placement of the anode (+) and cathode (-) along the nerve. **(C)** Dynamic gastric MRI images were collected before and during VNS. Every VNS setting was performed for 1 minute followed by another minute of rest. The sequence of different VNS settings was randomized within and across animals. **(D)** Example images obtained from dynamic gastric MRI.

### MRI Data Acquisition

Dynamic gastric MRI images were acquired with a 7-Tesla horizontal-bore small-animal system (BioSpec 70/30, Bruker, Billerica, MA, USA) and an imaging protocol adopted from our previous study^20^. Briefly, an abdominal localizer was first applied to reveal the long axis of the stomach from T2-weighted sagittal images. Then, a Fast Low Angle Shot gradient echo (FLASH) sequence with 4 slices was prescribed along the long axis of the stomach. The 4 slices were carefully positioned and adjusted to cover the antrum, pylorus and the duodenum (Fig. 1D). The MRI scans were acquired with TR/TE = 11.78/1.09ms, FA = 25°, 4 oblique slices, slice thickness = 1.5mm, FOV = 60 × 60 mm^2^, in-plane resolution = 0.4688 × 0.4688 mm^2^, and no averaging. To reduce motion artifact in the images, the MRI scans were respiratory-gated such that images were acquired only during the end-expiratory phase with minimum diaphragmatic motion. Readout gradient triggers were recorded to adjust acquisition times between volumes due to varying respiratory rates. The resulting sampling rate was typically between 2 to 3 seconds per volume, depending on the respiratory pattern of the animal.

### Experimental Protocol

After obtaining about 5 minutes of stable, baseline dynamic MRI images, cervical VNS was delivered simultaneously with MRI acquisition. The rats were allocated into 2 groups as follows: (i) a group of rats (N=8) that received afferent nerve stimulation (i.e. action potential propagates rostrally to the brain) and (ii) a group of rats (N=5) that received efferent nerve stimulation (i.e. action potential propagates caudally to end organs). The directionality of stimulation was biased by the placements of anode and cathode along the nerve. According to the principle of anodal blocking effect^22^, the nerve tissue at the cathode depolarizes and evokes compound nerve action potential (CNAP), while the nerve tissue at the anode hyperpolarizes and may block the propagation of CNAP. Therefore, as illustrated in Fig. 1B, placing the cathode rostrally to the anode and passing rectangular pulses of current between the two electrodes was biased for afferent nerve activation (hereinafter referred to as afferent VNS). Meanwhile, placing the cathode caudally to the anode may favor efferent nerve activation (hereinafter referred to as efferent VNS). In addition to evaluating the effect of afferent versus efferent VNS on gastric motility, we further applied bidirectional VNS (i.e. combined afferent and efferent VNS) to the rats in the afferent VNS group, immediately after which afferent VNS was performed. The bidirectional VNS was achieved by delivering rectangular pulses of current in alternating directions between the two electrodes as elaborated elsewhere^21^.

For each VNS group, the stimulus parameters were varied in terms of pulse amplitude (PA: 0.13, 0.25, 0.5, 1mA), width (PW: 0.13, 0.25, 0.5ms), and frequency (PW: 5, 10Hz). The frequency of afferent or efferent VNS was defined as the number of electrical pulses delivered per second. The frequency of bidirectional VNS was defined as the number of paired cathodal and anodal pulses delivered per second. Low, medium, and high values were selected for each parameter settings, all of which are frequently used in clinical and preclinical settings. Here, a stimulus dose was defined as the area under individual electrical pulse [product of pulse amplitude (mA) and pulse width (ms)], representing the electrical charge [Q; micro coulomb (μC)] delivered to the nerve tissue. For bidirectional VNS, the stimulus dose for a pair of cathodal and anodal pulses was defined as the charge under the cathodal pulse, because the effect of afferent versus efferent VNS was treated independently. As illustrated in Fig. 1C, each stimulation setting comprised of a duty cycle of 1-minute ON and 1-minute OFF, and different VNS settings were performed in a randomized order to eliminate any causal effect of one setting on the other. No stimulus was delivered during the OFF period. As a result, every rat in each group underwent VNS with 24 different sets of parameters.

### Image Analysis

The gastric antrum, pylorus, and the proximal duodenum were segmented from the lumen-enhanced MRI images by using a custom-built pipeline in Matlab (Mathworks, Massachusetts, USA) developed in our previous work^20^. Gastric volumes obtained at different times were rigidly co-registered to the first volume to minimize motion-induced displacements. To quantify antral motility, 16 cross-sectional planes that are perpendicular to the antral axis were defined at various locations (from 0 to 7.5mm distant to pylorus) along the axis. The segmented voxels within each cross-sectional plane were summed over all slices to quantify the cross-sectional area (CSA) of the lumen. As illustrated in Fig. 2A, a time series that represents the temporal change of CSA was obtained at every location along the antral axis (herein referred to the contraction time series). When an occlusive contraction wave arrived at a cross-sectional plane, a minimal CSA was attained. Due to irregular sampling caused by respiratory gating, the contraction time series were further resampled to 0.5Hz. For almost every rat studied in this work, occlusive contractions only occurred along the less curvature, whereas the greater curvature only exhibited non-occlusive contractions.

**Figure 2:**
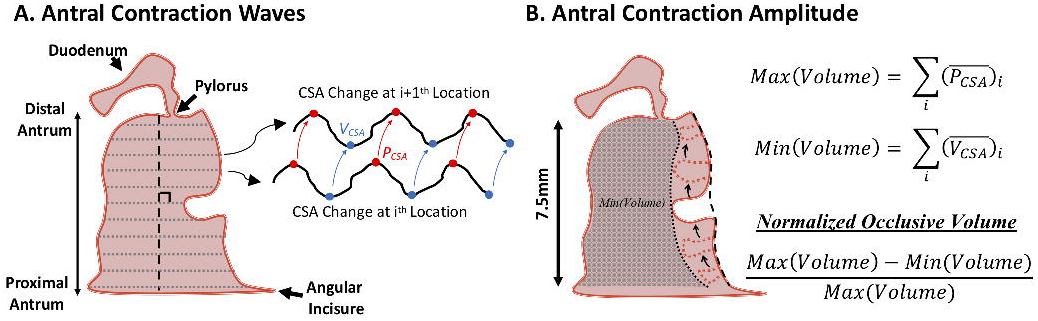
Image analysis of antral motility. **(A)** Cross-sectional areas (CSAs) along the long axis of the antrum (0 to 7.5mm distant from the pylorus, which correspond to 16 cross-sectional planes) were computed from the 3-D antral volume. The temporal change of CSAs at different locations of the antrum exhibits phase differences, revealing the propagation of antral contraction waves. When an occlusive contraction wave arrived at a cross-sectional plane, the minimal CSA was attained (V_CSA_). On the other hand, when the occlusive contraction wave was fully away from a cross-sectional plane, the maximal CSA was attained (P_CSA_) **(B)** Calculation of antral contraction amplitude. The contraction amplitude was defined as the normalized difference between the maximum and minimum antral volume, which represents the percentage occlusion of the antral volume. The maximum antral volume was defined as the sum of the temporal mean of peak values over all 16 cross-sectional planes, whereas the minimum antral volume was defined as the sum of the temporal mean of valley values over the 16 cross-sectional planes.

To compute the frequency of antral contraction waves, Fourier analysis was applied to all 16 contraction time series to derive each of their power spectral density (PSD). After averaging the 16 PSDs, the contraction frequency could be identified from the largest peak in the spectrum. To quantify the amplitude of antral contraction waves, a volumetric approach was proposed as shown in Fig. 2B. During the 1-min interval of VNS application, the peaks and valleys from all contraction time series were automatically detected. The peaks and valleys correspond to the maximal and minimal CSA of a cross-sectional plane, respectively. The maximum antral volume was then defined as the sum of the temporal mean of peak values over all 16 cross-sectional planes, whereas the minimum antral volume was defined as the sum of the temporal mean of valley values over the 16 cross-sectional planes. Finally, the percentage occlusion of the antral volume (herein referred to the antral contraction amplitude) was defined as the normalized difference between the maximum and minimum antral volume.

Next, pyloric motility was quantified as the area under the curve (AUC) of changes in luminal CSA at the pyloric sphincter. To compute luminal CSA at the pyloric sphincter, a distance transform-based approach was applied^23^. Specifically, distance transformation was first applied to the binary, segmented gut images. The distance transform map describes the distance from every luminal voxel to their closest boundary. Next, the medial axis of the GI tract was computed by applying a thinning algorithm (Matlab *bwmorph* function) that iteratively eroded pixels from the boundary until no more thinning was possible. Here, the distance-transformed values along the medial axis are the local maxima that represent the radius of the maximum inscribed circle, as illustrated in Fig. 3A. The cross-sectional diameter could be calculated and thus the luminal CSA was approximated by summing the product of diameter and slice thickness over all slices. Lastly, an integral analysis was applied to the contraction time series at the pylorus to obtain the AUC, as illustrated in Fig. 3B. Similarly, the contraction time series were resampled to 0.5Hz for subsequent frequency and amplitude analyses.

**Figure 3:**
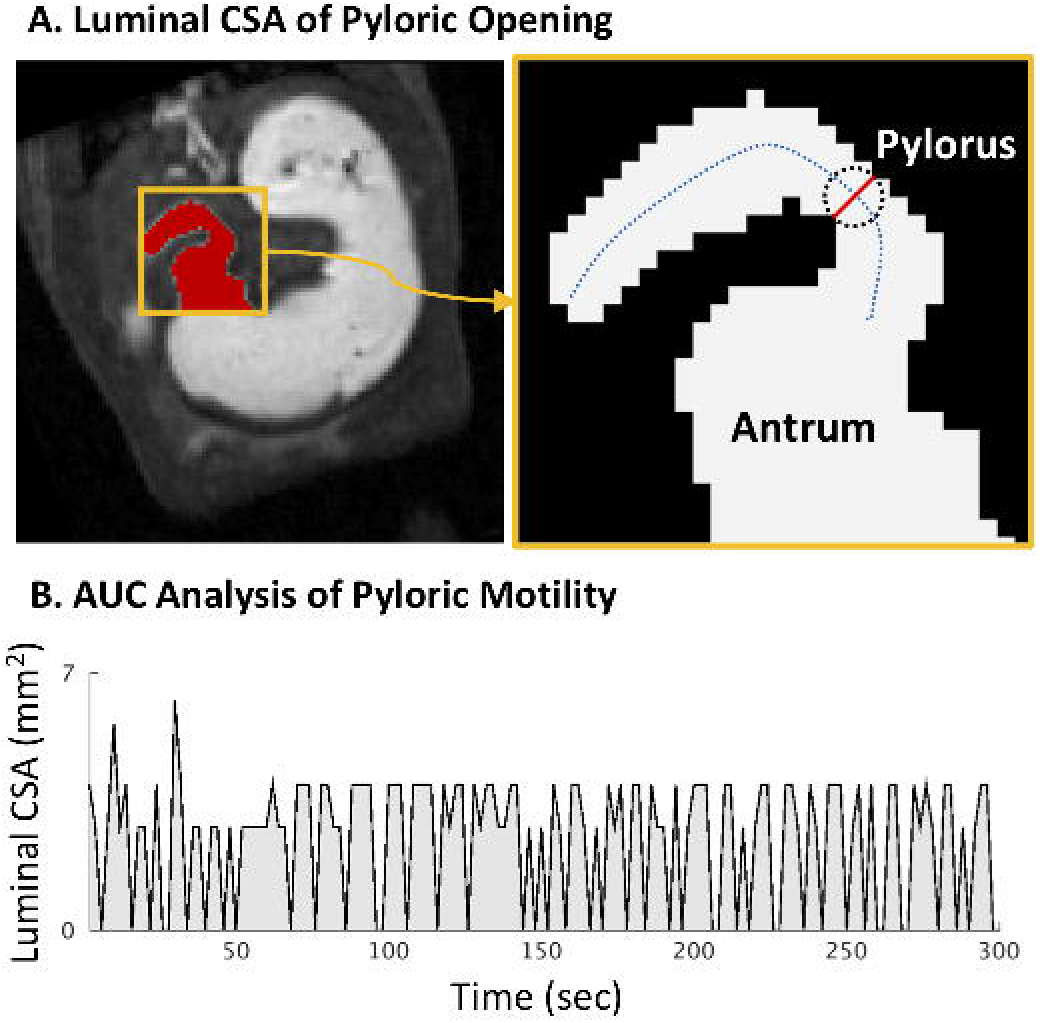
Image analysis of pyloric motility. **(A)** After the antropyloroduodenal region was segmented, the medial axis (blue dashed line) of the GI tract was computed, and the radius of the maximum inscribed circle (black dashed circle) along the medial axis was calculated from the distance-transformed binary segmented GI tract. The luminal CSA at the pylorus was quantified by summing the product of cross-sectional diameter (red line) and slice thickness over all slices. **(B)** The pyloric motility was defined as the area under the curve (AUC) of changes in luminal CSA of pyloric opening.

### Data Analysis

For each stimulation group, antral motility (i.e. contraction amplitude) and pyloric motility (i.e. AUC) were computed for each stimulus parameter setting during the 1-min ON period. The motility measurements were normalized against their baseline values over the 3-5 minutes immediately preceding the onset of the first VNS setting. The results were expressed as percentage increase/decrease from baseline. Further, a linear regression analysis was performed to characterize the association between stimulus dose and gastric motility. The independent variable used for the regression model was the logarithm-transformed stimulus dose, and the dependent variable was the mean antral contraction amplitude change or mean pyloric motility change. The linear regression analysis was conducted separately for the two stimulation frequencies (i.e. 5Hz and 10Hz).

### Statistics

All data are expressed as mean (SEM). A paired-sample Student’s t-test was conducted to compare motility measurements during each VNS setting to the same variable obtained during the baseline period. A Student’s t-test was applied to the slope obtained from regression analysis to test the goodness of fit of the regression model. Pearson correlations between the stimulus dose and the motility measurements were calculated as well. Values of p<0.05 were considered statistically significant.

## Results

We utilized contrast-enhanced MRI to investigate the effect of left cervical VNS on gastric motility in rats under a wide range of stimulus parameters. The effect of afferent versus efferent stimulation was dissociated, if not entirely, by configuring the placement of the cathode (i.e. the polarity) on the bipolar electrode. The combined effect of afferent and efferent VNS was further ascertained by delivering electrical pulses with alternating polarity to the vagus. Each VNS setting was delivered for one minute, followed by another minute of rest without stimulation, during which dynamic gastric MRI was performed continuously. The order of VNS settings was randomized within and across animals to counterbalance any causal effects. The occlusive amplitude of antral contraction waves and the tone of pyloric opening were measured as a function of stimulus settings. No significant change was observed in antral and pyloric contraction frequency across conditions. Table 1 shows the baseline motility measurements and Table 2 shows the VNS-evoked motility changes for each set of VNS parameters. Note that 1 rat from the afferent VNS group was excluded due to degraded image quality caused by abnormal respiratory pattern during imaging.

**Table 1.** Mean motility measurements in rats under baseline period. Data are presented in Mean (SEM).

| Group | Antral Motility |  |  | Pyloric Motility |  |
| --- | --- | --- | --- | --- | --- |
|  | Frequency (CPM) | Amplitude (% Occlusion) | Velocity (mm/s) | Frequency (CPM) | AUC (mm² · s) |
| Afferent VNS (N=7) | 4.46 (0.06) | 17.0 (0.8) | 0.77 (0.02) | 4.86 (0.25) | 0.59 (0.04) |
| Efferent VNS (N=5) | 5.25 (0.10) | 21.1 (2.0) | 0.85 (0.03) | 5.27 (0.18) | 0.74 (0.05) |

**Table 2.**
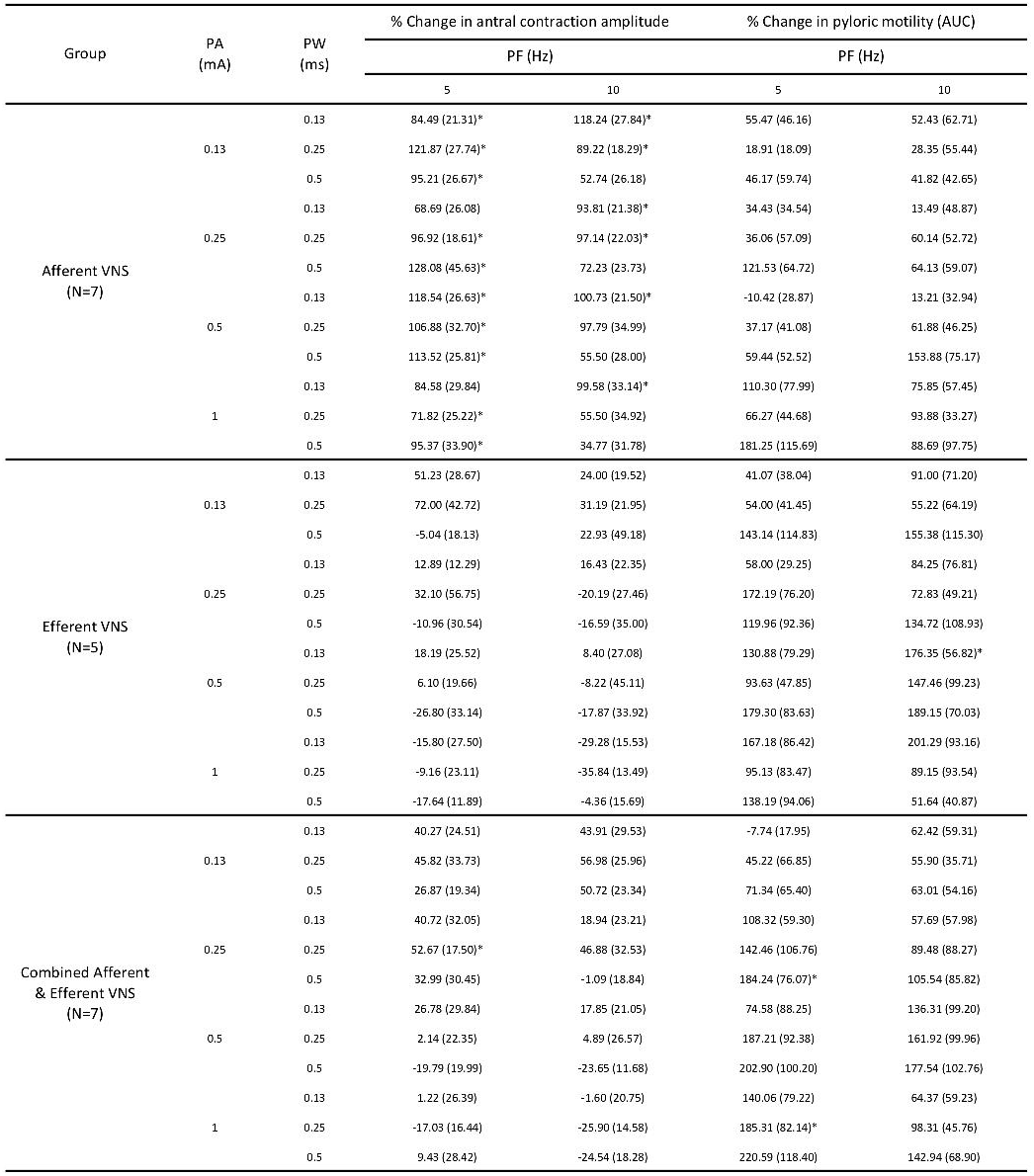
Mean percentage change in motility for each set of VNS parameters for the 3 conditions. Data are presented in Mean (SEM). An asterisk indicates a significant value (p<0.05).

### VNS effects on antral motility

For afferent VNS, the 24 sets of parameters primarily induced excitatory effect on antral contraction amplitude to various degree (Fig. 4A). At a stimulation frequency of 5Hz, there was a significant, positive change in contraction amplitude from baseline values under most stimulus settings. A non-significant increase (p=0.80) in contraction amplitude was found when increasing the stimulus dose (Q: product of pulse amplitude and pulse width) at this frequency. However, at the stimulus frequency of 10Hz, the contraction amplitude continued to fall as the pulse width was increased up to 0.5ms. Under this circumstance, an increase of VNS dose induced a significant (t=-3.54, p<0.01, R^2^=0.56) decrease in contraction amplitude, but the amplitude was overall still greater than baseline values.

**Figure 4:**
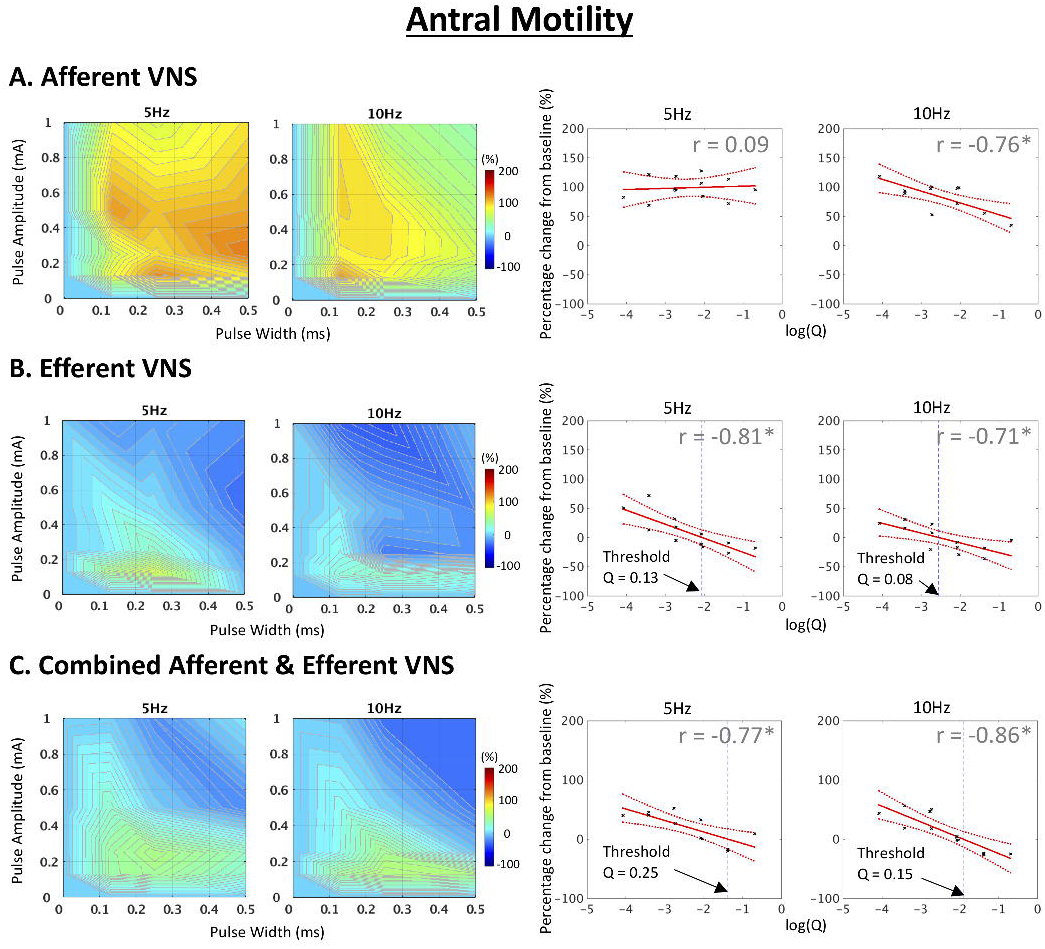
Effects of VNS parameters and directionality of VNS on antral motility. **(A)** Afferent VNS. **(B)** Efferent VNS. **(C)** Combined Afferent and Efferent VNS. **Left panel:** Effect of VNS on antral contraction amplitude under different VNS parameters. **Right panel:** Linear regression analysis of changes in antral contraction amplitude as a function of log-transformed VNS dose (Q: pulse amplitude x pulse width). Afferent VNS promoted antral contraction amplitude more effectively than efferent VNS. Meanwhile, high dose efferent VNS inhibited antral contraction. For both efferent and combined afferent and efferent VNS, a cut-off value Q (dotted blue line) was defined when the regression line crossed the zero-percentage change. The cut-off was higher for the stimulus frequency at 5Hz than at 10Hz. The color bar indicates percentage change of antral contraction amplitude from baseline values. Dotted red lines: 95% confidence interval of regression. r: Pearson correlation coefficient. *p<0.05.

Efferent VNS imposed two different effects on antral motility: one being excitatory and the other being inhibitory, as illustrated in Fig. 4B. At both stimulus frequencies, lower doses of VNS promoted contraction amplitude, with 5Hz being more pronounced than 10Hz. However, the results from linear regression (t=-4.08, p<0.01, R^2^=0.63 for 5Hz; t=-3.11, p<0.05, R^2^=0.49 for 10Hz) indicated a significant, proportional decrease in contraction amplitude as the stimulus dose increased, highlighting the recruitment of inhibitory fibers in the efferent pathway. A cut-off value Q was defined when the regression line crossed the zero-percentage change. The cut-off was higher for the stimulus frequency at 5Hz (Q=0.13μC) than at 10Hz (Q=0.08μC), suggesting that the threshold for activating inhibitory fibers was lower at 10Hz than at 5Hz.

Interestingly, when afferent and efferent VNS were performed alternately, the stimulus-response relations show an additive effect of afferent and efferent VNS, with the effect of efferent VNS being more dominant over afferent VNS (Fig. 4C). Combined afferent and efferent VNS could promote antral contraction amplitude under a wider range of low dose VNS settings than by performing efferent VNS alone. Regression analysis revealed a significant decrease in contraction amplitude as the stimulus dose increased (t=-3.51, p<0.01, R^2^=0.55 for 5Hz; t=-4.80, p<0.001, R^2^=0.70 for 10Hz). Similarly, a higher cut-off Q was found for the stimulus frequency at 5Hz (Q=0.25μC) than at 10Hz (Q=0.15μC).

### Effects of strong VNS dose on gastric physiology

As revealed in Fig. 4, we found that high dose efferent (as well as combined afferent & efferent VNS) dampened antral contraction amplitude. To further demonstrate the effect, we additionally performed high-dose efferent VNS (1mA, 0.5ms, 10Hz) with 30 seconds on, 30 seconds off duty cycle in one animal. As illustrated in Fig. 5A, when efferent VNS at supramaximal intensity was applied to the left cervical vagus nerve, gastric secretion was induced, resulting in an enlarged corporal and antral volume. Furthermore, antral contraction waves ceased during the ON period, highlighting the inhibitory effect of efferent VNS. Fig. 5B shows an example CSA change extracted from middle antrum before, during and after efferent VNS. At the onset of each VNS, an abrupt change in CSA can be observed, reflecting the enlargement of antral volume caused by relaxation of the antrum and the infused secretory volume. The propagation of antral contraction waves ceased during efferent VNS. Such inhibitory effect on antral contraction was immediately followed by intensive, rebound contractions when the VNS was turned off. The effect was reproducible over multiple cycles. Supplementary video 1 demonstrates the dynamic change in antral contraction waves during high dose efferent VNS.

**Figure 5:**
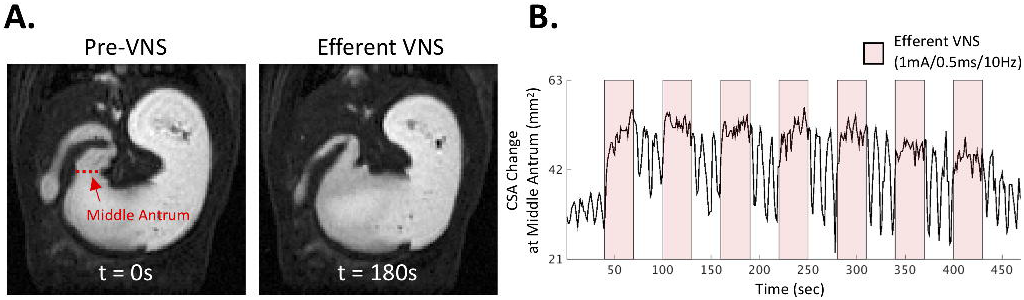
High dose efferent VNS retards antral peristalsis and induces gastric secretion. **(A)** Example gastric images before and during high dose efferent VNS (PA=1mA, PW=0.5ms, PF = 10Hz). Efferent VNS under this parameter setting ceased antral contraction wave while inducing excessive secretory volume into the corpus and the antrum. A dynamic illustration of this process can be found in supplementary video 1. **(B)** A CSA change taken from the middle antrum as indicated in panel **(A)**. The onset of high dose efferent VNS significantly increased the CSA, followed by little or no phasic change in CSA. There was a large rebound, intensive contraction on the offset of efferent VNS.

### VNS effects on pyloric motility

Regardless of the directionality of VNS, the vagal innervation on pyloric sphincter was found to be mostly relaxatory (corresponding to a decrease in pyloric tone, or equivalently to an increase in AUC). For afferent VNS at 5Hz, the maximum response was induced either at the highest PA (1mA) or the highest PW (0.5ms) utilized in this study. When stimulating at 10Hz, the greatest response was observed at PA=0.5mA and PW=0.5ms, and the effective settings for promoting pyloric motility were of wider range than that of 5Hz. An increase in stimulus dose induced a significant increase in pyloric AUC at both stimulus frequencies (t=2.30, p<0.05, R^2^=0.35 for 5Hz; t=3.02, p<0.05, R^2^=0.50 for 10Hz), as shown in Fig. 6A.

**Figure 6:**
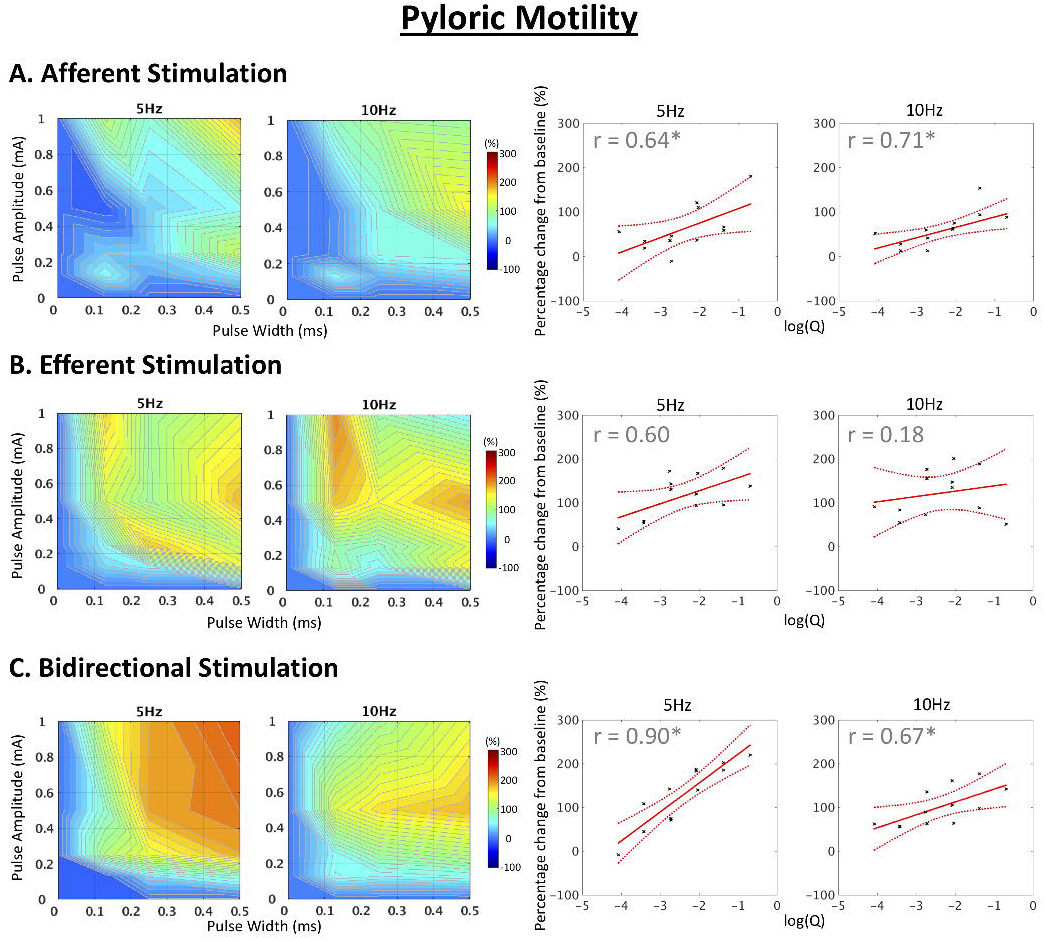
Effects of VNS parameters and directionality of VNS on pyloric motility. **(A)** Afferent VNS. **(B)** Efferent VNS. **(C)** Combined Afferent and Efferent VNS. **Left panel:** Effect of VNS on pyloric motility under different VNS parameters. **Right panel:** Linear regression analysis of changes in pyloric motility as a function of log-transformed VNS dose (Q: pulse amplitude x pulse width). An increase in stimulus dose generally promoted pyloric motility, indicating an increase in AUC of pyloric opening (or equivalently a decrease in pyloric tone). The color bar indicates percentage change of AUC of pyloric opening from baseline values. Dotted red lines: 95% confidence interval of regression. r: Pearson correlation coefficient. *p<0.05.

On the other hand, efferent VNS was found to promote pyloric opening more effectively under a wider spectrum of VNS settings than afferent VNS (Fig. 6B). This effect once again elaborates the activation of inhibitory fibers in the efferent pathway. However, depending on the strength of the stimulus, efferent VNS induced a marginally significant increase and a slight, non-significant increase in pyloric AUC for 5Hz and 10Hz, respectively (t=2.16, p=0.06, R^2^=0.32 for 5Hz; t=0.66, p=0.52, R^2^=0.04 for 10Hz).

A marked increase in pyloric motility was found when performing combined afferent and efferent VNS at high dose and at 5Hz (Fig. 6C). The stimulus-response relations again show additive effect of afferent and efferent VNS. A stimulus frequency of 5Hz was found to promote pyloric motility more effectively than 10Hz. Finally, a significant linear regression indicated a proportional increase in pyloric motility as the stimulus dose increased (t=6.26, p<0.001, R^2^=0.80 for 5Hz; t=2.59, p<0.05, R^2^=0.40 for 10Hz).

## Discussion

In this study, we characterized the effect of different cervical VNS settings on gastric motility in anesthetized rats. To our knowledge, this is the first animal study that utilizes gastric MRI to non-invasively assess postprandial gastric motility under VNS with a wide range of parameters. The impact of directionality of VNS was studied by configuring the cathode on the bipolar cuff electrode. Gastric MRI revealed that electrical activation of the afferent pathway may promote antropyloric motility more effectively then directly activating the efferent pathway. A reduction in antral contraction amplitude and relaxation of pyloric sphincter under efferent VNS highlighted the inhibitory pathway of the motor limb of the vagovagal circuitry.

### Efferent VNS induces multiple effects on gastric physiology

Previous studies on vagal control of gastric motility have largely focused on the efferent parasympathetic innervation of the GI tract^10,12,24,25^. The modulatory role of efferent VNS with different stimulus parameters on gastric motility was discovered by Veach^26^ over a hundred years ago. In a cat model, he found that the stronger the stimulus strength and/or the higher the stimulus frequency, the more prominent the inhibitory effect on gastric motility. The present finding supports the general opinion that there are two groups of efferent fibers that influence gastric motility: one being excitatory and the other being inhibitory^2^. In line with previous studies^10,27^, the activation threshold for the two groups of fibers seems to be different, which was dependent on the stimulus strength (pulse amplitude and pulse width) as well as the rate at which these pulses were applied (frequency). Here, we found that lower pulse amplitude and/or lower pulse width produced excitatory response on antral contraction amplitude. However, when we continued to increase the pulse amplitude and pulse width over a certain level, the amplitude of antral contraction waves became smaller and eventually, the contraction waves ceased at supramaximal intensity (Supplementary video 1). The phenomenon confirms the presence of efferent fibers with two different calibers; the fiber caliber for inducing excitatory response is perhaps larger than the fiber caliber for inducing inhibitory response, where the activation threshold is lower for the former than the latter. The two fiber groups are likely unmyelinated fibers, as the activation thresholds for the two groups were suggested to be both higher than that needed for inducing cardiac response^28,29^, under which the heart is primarily mediated by efferent B-fibers. Our experimental observation is also consistent with the finding that there are only a few myelinated fibers in the abdominal vagi^30^. Notably, the pulse amplitude and pulse width required for inducing inhibitory response are generally smaller than previously reported parameters, though different animal species were used^10,12^. Speculatively, the difference in experimental condition (i.e. pre-prandial versus post-prandial state) may be the underlying cause, as the basal gastric tone should vary across the two states. Interestingly, the threshold for producing inhibitory response on antral motility was slightly higher at 5Hz than at 10Hz. It was suggested that the stimulus frequency could be responsible for selective activation of the efferent excitatory (cholinergic) and inhibitory [non-adrenergic, non-cholinergic (NANC)] effects^31^ (i.e. 5Hz for acetylcholine release and 10Hz for vasoactive intestinal peptide release). Nevertheless, a clear cut on frequency-specificity was not apparent in the present study, as the 2 frequencies both produced excitatory and inhibitory response.

The excessive secretion induced by efferent VNS observed in the present study is in complete agreement with previous findings^24,32–34^. The effect was most visually pronounced at the supramaximal intensity, regardless of the 2 stimulus frequency utilized in this study. This finding is perhaps not too surprising, as gastric parietal cells in the rat are mainly innervated by unmyelinated C-fibers^33,35^, therefore a larger stimulus dose is required to trigger vagally mediated secretory response. On the other hand, the inhibitory motility pattern induced by high dose efferent VNS consists of two phases. The first phase was a rapid relaxation of the antrum at the onset of stimulus, followed by intensive, rebound contractions on cessation of the stimulation. As reported and discussed elsewhere^12,36^, the rebound contractions could be of purinergic origin and the release of prostaglandins by NANC terminals may account for this poststimulus pattern.

The influence of efferent VNS on pyloric motility appeared to be primarily inhibitory. This observation coincides with previous findings that VNS does not seem to impose a strong tonic influence on pyloric resistance; VNS was shown to decrease pyloric resistance through the NANC pathways and thus increase transpyloric flow^37^. However, the seemingly pure inhibitory effect could depend on the choice of stimulus frequency, as it was suggested in a dog study that low-frequency stimulation (0.2-0.5Hz) exerted tonic contractions, whereas higher frequency (>0.7 Hz) stimulation could inhibit both phasic and tonic contractions^38^. Further experiments are needed to evaluate the potentially excitatory effect of efferent VNS on pyloric tone with a wider range of stimulus frequency.

In sum, our results suggest that stimulating the vagal efferent nerves could induce multiple effects on gastric physiology. These effects were heavily, if not entirely, dependent on the dose and the frequency of the stimulus. The likelihood that both excitatory and inhibitory efferent fibers are unmyelinated fibers pose technical difficulty in selective activation of one but not the other fiber group. As a result, efferent VNS employ complex and heterogeneous neural signaling to the gut, which may lead to undesired, mixed gastric responses. Furthermore, a promotion of antropyloric coordination seems difficult to achieve, because there exists no single parameter setting that can optimally increase antral motility while reducing pyloric tone.

### Reflex excitation on gastric motility via afferent VNS

While most existing studies utilized electrical stimulation of the peripheral end of the sectioned vagal nerve, the effect of afferent VNS on gastric motility has received considerably less attention. Here, we performed afferent VNS at the cervical level to mimic the vagovagal reflex occurring in the physiological state. The vagal afferents carry signals from stretch receptors^39,40^ and chemoreceptors^41^ to dorsal vagal complex in the brainstem, resulting in downstream, coordinated signals to different segments of the GI tract^2,42^. It has been widely demonstrated that fundic distention could induce reflex excitation on antral motility^12,14^ and pyloric relaxation^43^. On the other hand, balloon distension of the duodenum could inhibit antral motility and increase pyloric tone^44,45^. Speculatively, activation of different branches of the vagal afferents may result in selective and choreographic gastric motility. Indeed, we found that afferent VNS at 5Hz significantly promoted antral motility under a wide range of stimulus settings. This finding suggests the afferent fibers that are responsible for delivering excitatory response to the antrum are recruited under this stimulus frequency. When the stimulus frequency was increased to 10Hz, a higher stimulus dose resulted in a decrease in contraction amplitude, suggesting that a second group of fibers has been recruited. The second vagal afferents may convey reflex signals that return to the stomach to activate relaxation of the smooth muscle via inhibitory vagal efferents, as similar finding was reported previously^46,47^. Such relaxatory response is also consistent with our finding that a decrease in pyloric tone could be achieved at a similar stimulus dose. Taken together, electrical activation of the vagal afferent pathway into the central nervous system could potentially result in a more coordinated, physiological downstream signaling to different segments of the GI tract than direct efferent VNS.

### Limitations and future directions

There are several limitations in this study. Firstly, the directionality of VNS was controlled by the placement of the cathode on the bipolar cuff electrode. This approach relies on the phenomenon known as “anodal block”, where the anode should be theoretically hyperpolarized and thus blocking the propagating CNAP evoked at the cathode. However, whether the evoked CNAP was indeed unidirectional was not confirmed in all rats. Although the stimulus-response maps show differential patterns between conditions within (afferent vs. combined afferent & efferent) and across (afferent vs. efferent) animals, we cannot rule out the possibility that the stimulation was not fully unidirectional. Additional experimentation (e.g. vagotomy, chemical nerve blockade, etc.) is required to affirm the directionality of VNS.

Secondly, all animals were sedated with dexmedetomidine and low-dose Isoflurane (<0.5%). The use of anesthetics could have impacted the excitability of the nerve and thus might have modulated gastric motility. Dexmedetomidine, also known as an agonist of α2-adrenergic receptors, has an inhibitory effect on the sympathetic system^48^. While the sympathetic nervous system primarily plays an inhibitory role on gastric motility^1^, reducing sympathetic innervations on the GI tract may potentially alter the pattern of VNS-induced gastric responses. Further experiments may be of interest to evaluate the VNS effect on animals under different anesthetics.

Last but not least, a majority of VNS-gastric studies were conducted on fasted animals. The pattern of pre-prandial gastric physiology is relatively more stable and predictable. However, postprandial gastric motility is a dynamic process that includes complex feedforward, feedback and local reflexes during digestion of a meal. It is therefore not unlikely that the VNS-induced gastric response observed in the present study might be biased by spontaneous changes from the gastric emptying process. Although the sequence of the 24 VNS settings was randomized across animals so that the impact of spontaneous gastric activity could be counterbalanced, it remains an experimental challenge to screen the effects of various VNS parameters on the same animal within a single imaging session.

## Acknowledgement

This work was supported in part by NIH SPARC 1OT2TR001965 and Purdue University.

---

**Supplementary Video 1. Strong efferent VNS retarded antral peristalsis and induced gastric secretion.** Efferent VNS was delivered at 1mA, 0.5ms and 10Hz, with a duty cycle of a 30-second ON period followed by 30 seconds of rest. The video was fast-forwarded and displayed at a frame rate of 9 volumes second^−1^.

